# Characterisation of *DMPK* and MBNL1 expression in cell models of Myotonic Dystrophy: A platform for drug screening

**DOI:** 10.1101/2025.01.31.632773

**Authors:** Andrea López-Martínez, Sergio Martín-González, Noemi Torres-Conde, Nahia Alcalá-Manso, Abdullah Al-Ani, Adolfo López de Munain, Anne Bigot, Kamel Mamchaoui, Gisela Nogales-Gadea, Virginia Arechavala-Gomeza

**Affiliations:** Nucleic Acid Therapeutics for Rare Disorders (NAT-RD), Biobizkaia Health Research Institute, Barakaldo, Spain; Group of Neurosciences, Departments of Pediatrics and Neuroscience, Faculty of Medicine and Nursing, University of Basque Country (UPV/EHU), 20014 Donostia-San Sebastian, Spain; Groups of Sensorial Neurodegeneration and Neuromuscular Diseases, Neuroscience Area, BioGipuzkoa-BioDonostia Health Research Institute (IIS Biodonostia), 20014 Donostia-San Sebastian, Spain; CIBERNED, ISCIII (CIBER, Carlos III Institute, Spanish Ministry of Sciences and Innovation), 28031, Madrid, Spain; Department of Neurology, Hospital Universitario Donostia. OSAKIDETZA, 20014 Donostia-San Sebastián, Spain; Department of Medicine, School of Medicine, University of Deusto, Bilbao, Spain; MyoLine, Sorbonne Université, Inserm, Institut de Myologie, Centre de Recherche en Myologie, F-75013 Paris, France; Grup de REcerca Neuromuscular de BAdalona (GRENBA), Institut d’Investigació en Ciències de la Salut Germans Trias i Pujol (IGTP), Campus Can Ruti, Universitat Autònoma de Barcelona, 08916 Badalona, Spain; Ikerbasque, Basque Foundation for Science, Bilbao, Spain

**Author notes:** **Corresponding Author:** Prof. Virginia Arechavala-Gomeza, Nucleic Acid Therapeutics for Rare Diseases, Biobizkaia Health Research Institute, Plaza de Cruces S/N, 48903 Barakaldo, Spain. **Author contributions:** Conceptualization, V.A.-G. Methodology, validation and investigation, A. L.-M.; S.M.-L.; N.T.-C.; N.A.-M.; A.A.-A.; V.A.-G. Formal analysis, A. L.-M.; V.A.-G. Resources, A.B.; K.M.; V.A.-G. Writing-original draft preparation A. L.- M. Writing-review and editing, A. L.-M.; S.M.-L.; A.A.-A.; A.L.-M.; G.N.-G.; V.A.-G. Visualisation, supervision, A. L.-M. V.A.-G; project administration and funding acquisition, V.A.-G. **Competing interest statement:** The authors declare no competing interests.

**Keywords:** Myotonic dystrophy, cell models, drug screening, antisense oligonucleotides, MBNL1, DMPK

## Abstract

Myotonic dystrophy type I (DM1) is caused by CTG repeat expansions in the *DMPK* gene leading to mRNA toxicity and sequestration of the splicing regulator MBNL1, affecting many tissues. We have developed an *in vitro* screening platform based on ddPCR and in-cell western to quantify these mRNAs and proteins and characterised more than 20 cell models to define DM1 biomarkers that could be useful for drug screening. DMPK protein levels were reduced in DM1-immortalised myoblasts and myotubes, but not in fibroblasts, while MBNL1 protein was consistently lower in all DM1 myogenic cultures, whether primary or immortalised. Myogenic differentiation of cultures led to an increase in *DMPK* mRNA expression, which was translated into increased MBNL1 sequestration in foci. We further corroborated the platform’s ability to assess therapeutic outcomes, evaluating the effect of a DMPK gapmer ASO and one siRNA: while the gapmer increased MBNL1 protein levels, the siRNA had no significant effect on MBNL1 release. Our platform and the in-depth characterisation of some of the most used models would be of use to the DM1 research community.

**Significance statement:** Myotonic dystrophy type I (DM1) is a multisystemic disease with a complex pathogenesis and multiple outcome measures for drug assessment *in vitro*. In the last years, the increasing number of new potential therapies targeting DM1 in clinical trials has increased the need for robust and rapid evaluation of preclinical candidates, as well as in-depth knowledge of the cell models used. Here, we present a new cell-based platform that enables robust quantification of DMPK and MBNL1 in cell culture for cell model characterisation and drug screening. Indeed, we highlight the differences observed in DMPK and MBNL1 protein quantification in primary fibroblasts and myotubes, immortalised fibroblasts, myoblasts and myotubes. We then, we performed a proof-of-concept drug evaluation of potential therapeutic strategies targeting DMPK, showing the most suitable for targeting the *DMPK* expanded transcript.

## Introduction

Myotonic dystrophy type I (DM1) is a complex autosomal disease caused by expansion of CTG trinucleotide repeats in the 3’ untranslated region of the *DMPK* gene. It is the most common form of adult muscular dystrophy, affecting 1 in 8000 people. DM1 is characterised by a variety of multisystemic symptoms caused by a complex molecular pathogenesis, including toxicity of *DMPK* mRNA transcripts and nuclear sequestration of MBNL1 protein, leading to deregulation of multiple signalling pathways (1–3). Various therapeutic strategies such as small molecules, antisense oligonucleotides (ASOs) and gene therapies, have been tested in DM1 *in vitro*, *in vivo,* and in a handful of clinical trials. Most therapies, through different mechanisms of action, aim to reduce RNA toxicity by decreasing *DMPK* mRNA transcripts or preventing MBNL1 binding to the expanded transcript (4).

DM1’s complex molecular pathogenesis is compounded by the large number of tissues affected by the disease. While numerous studies have been conducted to understand the molecular mechanism of the disease and to test therapeutic strategies, the limited availability of patient-derived cell lines poses a challenge to research. Fibroblasts, which are easily obtained from a skin biopsy, are preferred for cell culture studies over myoblasts, which require more invasive muscle biopsies. However, fibroblasts have not been extensively studied as DM1 cell models and further characterisation is needed to identify appropriate biomarkers (5–8).

Current methods for characterising cell models and assessing therapeutic response focus on quantifying foci and alternative splicing defects, but there is no consensus on the protocols for this assessment and results can vary between research groups. Although technical developments in recent years have allowed more automated analysis of foci and improved splicing analysis through transcriptomics, a simple and straightforward platform could accelerate treatment screening. Furthermore, it remains unclear how reducing foci or improving splicing may correlate with phenotypic improvements. While some therapeutic approaches have progressed to clinical trials, only a few patients have experienced symptomatic improvement. With many small molecules and antisense oligonucleotides (ASOs) available for further testing, rapid *in vitro* evaluation of different therapeutic approaches is essential for the development of effective treatments (9–13).

In light of all this, in this work we have optimised a platform based on In-Cell Western (ICW), a quantitative immunofluorescence assay that enables direct protein quantification in cell cultures, integrating the specificity of western blots with the efficiency and scalability of an ELISA. Indeed, it is a versatile technique with multiple applications in different fields (14) (15). In the field of neuromuscular disorders, our group previously developed a myoblast/myotube ICW (or myoblot) to quantify dystrophin (16) and myoblots have since been used to quantify dystrophin, utrophin and other proteins after exon skipping experiments in Duchenne muscular dystrophy cultures (17), and to characterise cell models newly generated by gene edition (18).Thus, we built on this expertise to quantify DMPK and MBNL1 expression in DM1 cell models, characterise these models, and identify useful biomarkers for *in vitro* drug screening. We complemented the protein validation of this platform with the absolute quantification of corresponding RNAs by digital droplet PCR. In this manuscript, we describe this platform, we use it to characterise a wide range of cell culture models, and we validate it further by testing two potential DM1 therapeutic strategies: a DMPK gapmer previously described (19) and a DMPK dsiRNA.

## Results

### Optimisation of In-Cell Western protein quantification

An In-Cell Western (ICW) assay is a quantitative immunofluorescence assay performed in 96-well plates, as in our case, or 384-well plates. Cells are seeded in a 96-well plate and fixed, permeabilised and stained in the same well. The protein is thus detected *in situ*, in its natural conformation and intracellular location, without the need for denaturing (Figure 1). Protein quantification by In-Cell western assays requires accurate cell seeding density, which needs to be within a linear range and permits the quantification of cell density in the same well as target protein. To define the linear range of detection of the cell number stain used for this purpose, we seeded different cell densities of control-derived immortalised myoblasts, and we stained with Cell Tag 700 stain (Supplementary Figure 1A).

**Figure 1.**
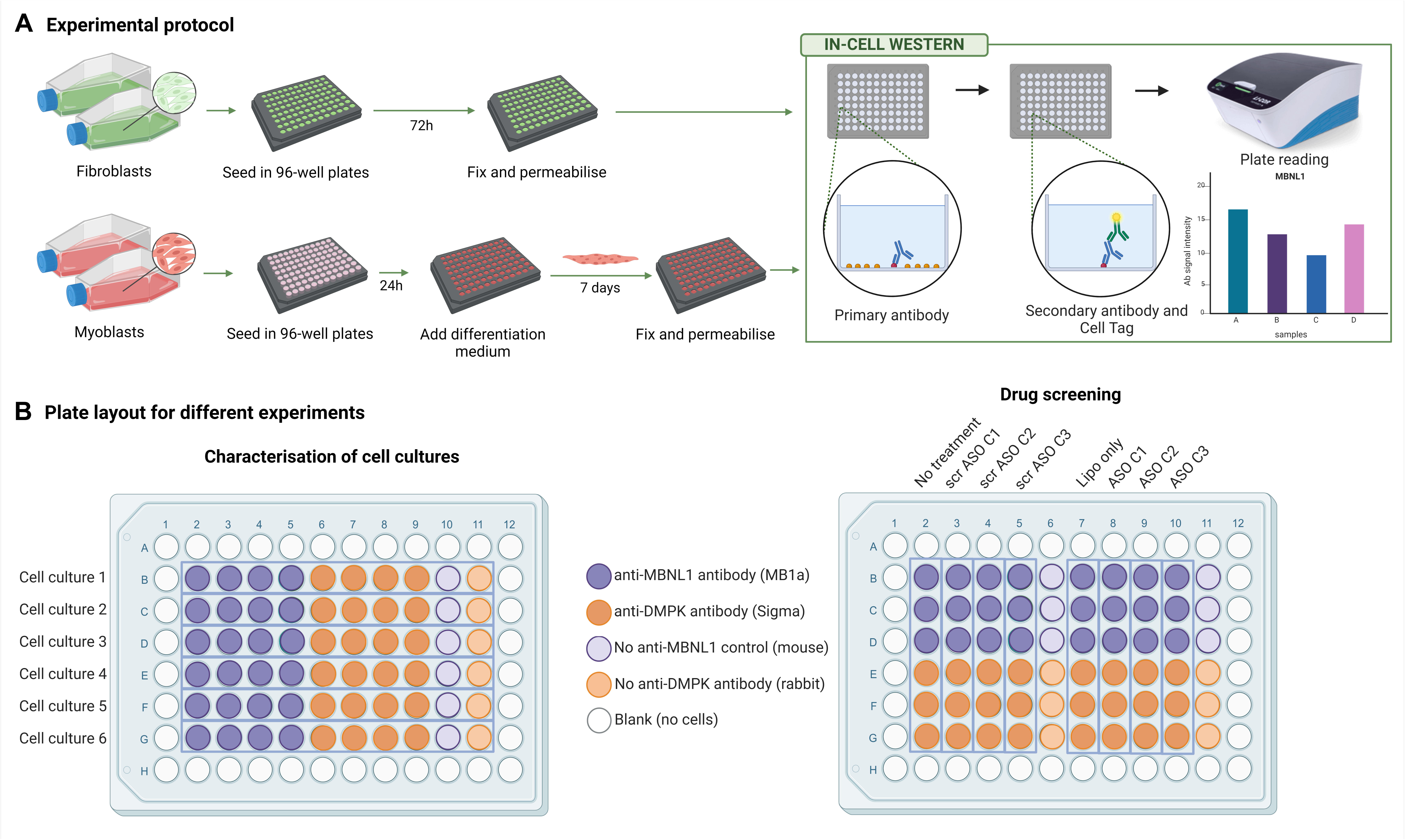
In-Cell western workflow and experiment design. **(A)** Schematic representation of cell culture and In-Cell western procedure for protein quantification in fibroblasts and myotubes. **(B)** Example of plate set-up for cell culture characterisation or ASO testing at three different concentrations (C1, C2, C3). No treatment refers to cells no subjected to transfection and lipo only refers to cell transfected without ASO (negative control).

As ICWs do not have the resolution of microscopy nor the possibility of confirming the size of the protein detected, the antibodies used need to be thoroughly validated.

While *DMPK* transcript levels have previously been assessed in DM1 to understand the role of *DMPK*-expanded transcripts in DM1 pathogenesis, fewer studies have focused on quantifying DMPK protein in cell culture (20, 21). Difficulties in assessing the role of DMPK protein implications in different features of the disease are due in part to the lack of validated antibodies (22). Several DMPK antibodies with different targeting epitopes are commercially available, but there is no consensus on which are the best ones for targeting DMPK in cell culture. Therefore, we tested different DMPK antibodies on control myotubes to perform a parallel validation of their targets. After discarding some antibodies for unspecific binding, we evaluated HPA007164 from Sigma and MANDM5 from the Monoclonal Antibody Database from the Wolfson Centre for Inherited Neuromuscular Disease (CIND). Both showed the expected values described in the literature for DMPK recognition: correct localisation under the microscope (cytoplasm, endoplasmic reticulum and mitochondria) (23) (Figure 2A), and a size, when evaluated by Jess simple western blotting, of 82kDa, consistent with the molecular weight reported in the literature (79kDa) (24) (Figure 2C). We selected the Sigma as it showed a more robust signal.

**Figure 2.**
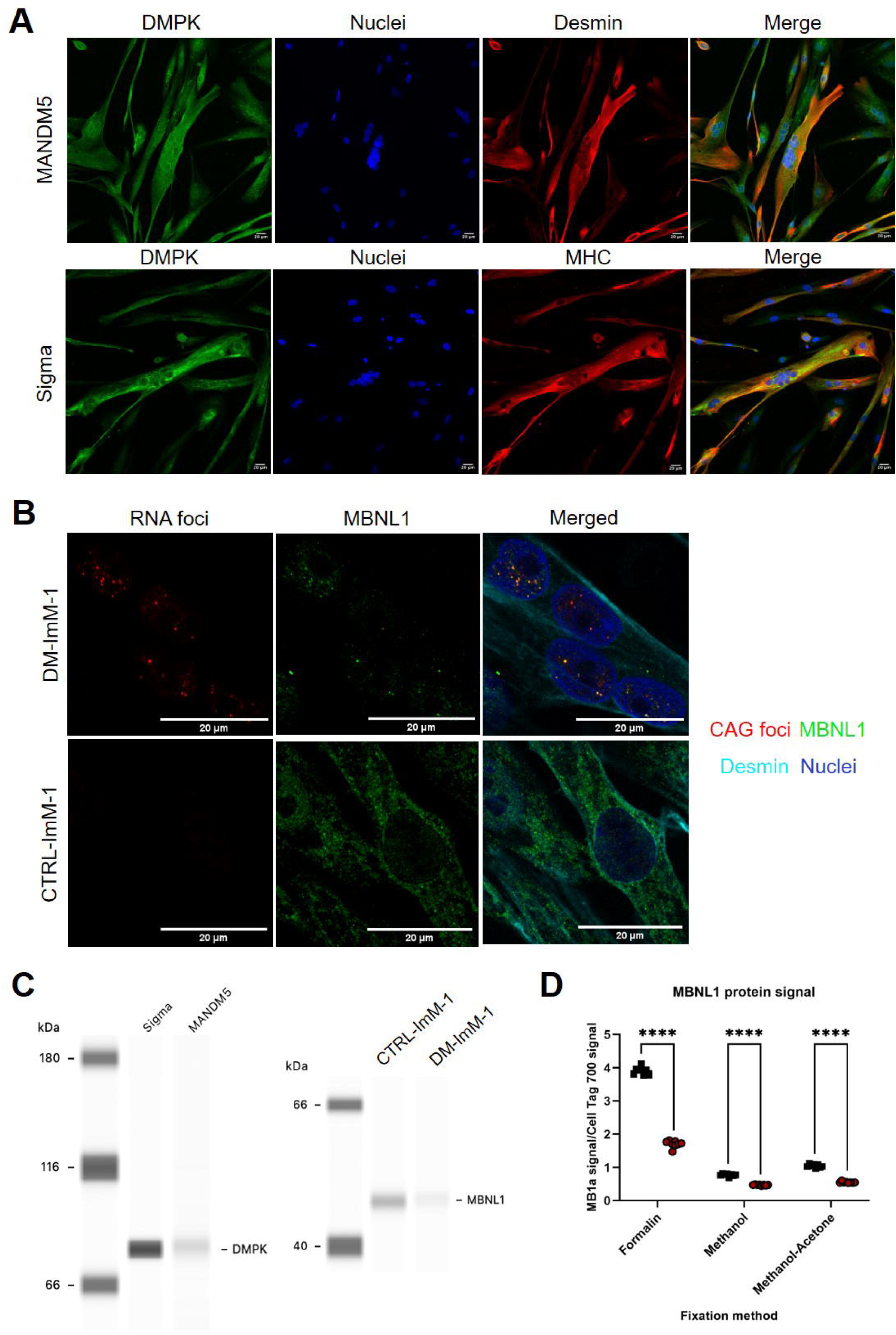
Antibody selection for DMPK and MBNL1 quantification. **(A)** Immunofluorescent staining of control immortalised myotubes with two DMPK-targeting antibodies: CTRL-ImM-1 myotubes differentiated for seven days were stained with HPA007164 (Sigma) and MANDM5 DMPK antibodies (green), nuclei (blue) and desmin (red) or MHC (red) showing the expected localisation. **(B)** Immunofluorescent staining of control (CTRL-ImM-1) and DM1 immortalised myotubes (DM-ImM-1) with MBNL1 antibody MB1a and (CAG)_7_ probe. CUG foci are seen as small red points in DM1 cultures, MBNL1 is stained green, desmin is cyan and nuclei are blue). **(C)** Representative image of Jess simple western lane visualization of DMPK-targeting antibodies in CTRL-ImM-1 and MB1 antibody targeting MBNL1 in CTRL-ImM-1 and DM-ImM-1 myoblasts. **(D)** MB1a normalised data by Cell Tag 700 stain signal in CTRL-ImM-1 and DM-ImM-1 immortalised myoblasts after fixation with different methods. Data is represented as mean ± SEM of 3 technical replicates per experiment in three independent experiments. Significant differences were determined by two-way ANOVA and Bonferroni’s multiple comparison test (**** p<0.0001).

The quantification of MBNL1 protein, on the other hand, has been of great interest in the development of drugs aimed at increasing *MBNL1* expression or releasing the protein from the *DMPK-*expanded transcripts. Detection of the MBNL1 protein signal is often performed by immunocytochemistry or western blot after treatment with potential therapeutic compounds to assess MBNL1 release from foci and to study treatment efficacy (25, 26). The most commonly used antibody, MB1a (clone 4A8) from the Wolfson Centre for Inherited Neuromuscular Diseases (CIND) monoclonal antibody database, targets an epitope encoded by exon 3 of the MBNL1 protein, which is constitutively expressed in all MBNL1 isoforms (27). We corroborated the validity of this antibody testing it on DM1 and control immortalised myotube by FISH and immunocytochemistry and Jess simple western, as represented in Figures 2B and C (respectively). It showed the expected distribution of MBNL1, co-localizing with RNA foci, which was also detected by a (CAG)_7_ probe in the nucleus (Figure 2B). When with this antibody was used to analyse by extracts of both control and DM1 cultures by Jess, it showed a band at the expected molecular weight, 48 kDa, (Figure 2C) (28).

As the entrapment of MBNL1 in foci is at the core of the main pathogenesis theories in DM1 and its release is the mechanism of several potential treatments, we were interested to know if we were able to measure free or “soluble” MBNL1 by ICW. ICW relies on standard immunohistochemical techniques but lack the resolution of confocal microscopies. Recently, it was suggested that fixation of DM1 and control cell lines with 4% PFA (paraformaldehyde) or 50% Methanol-50% Acetone could affect MBNL1 protein detection (29): methanol-acetone fixated cells show a preference for insoluble MBNL1 binding to *DMPK*-expanded transcripts, whereas PFA-fixed cells show a more dispersed pattern, corresponding to insoluble MBNL1, but also soluble MBNL1 in the nucleus, which is also present in control cell lines (29)(^29^). We, therefore, decided to test this hypothesis and compared MBNL1 distribution by ICW when DM1-ImM-1 and CTRL-ImM-1 immortalised myotube cultures where fixed with formalin, ice-cold methanol (Methanol) and 50% methanol-50% acetone (Methanol-Acetone) fixation methods (Figure 2D). The resolution of the ICW data, obtained by Odissey M plate scanner, is 100μm, which defines the distance by which the instrument distinguishes between two points of signal intensity. As shown in Figure 1B, the cell nucleus is approximately 20μm, suggesting that we are unable to distinguish different points within the nucleus, i.e. foci. The intensity of the foci is dispersed by the intensity of the surroundings, suggesting that the MBNL1 signal detected by ICW reflects the diffuse MBNL1 signal observed in control and DM1 myotubes and not the foci-specific signal from DM1 cultures. Quantification by ICW using the Odissey M scan shows statistically significant differences between control and DM1 samples for all fixation methods. However, the signal is higher in formalin-fixed wells compared to methanol and methanol-acetone wells, suggesting that we are detecting the diffuse MBNL1 signal previously observed after 4% PFA fixation (29). Lower levels of diffuse MBNL1 are observed when fixed with methanol and methanol-acetone, similar to methanol-acetone fixed cells in (29).

### Validation of the platform on immortalised control and patient cultures

Once the ICW methodology was optimised, it was combined with absolute RNA quantification by digital droplet PCR and this platform was used to characterise control and DM1 patient cultures described in (30) by assessing DMPK and MBNL1 expression at RNA and protein levels (Figure 3).

**Figure 3.**
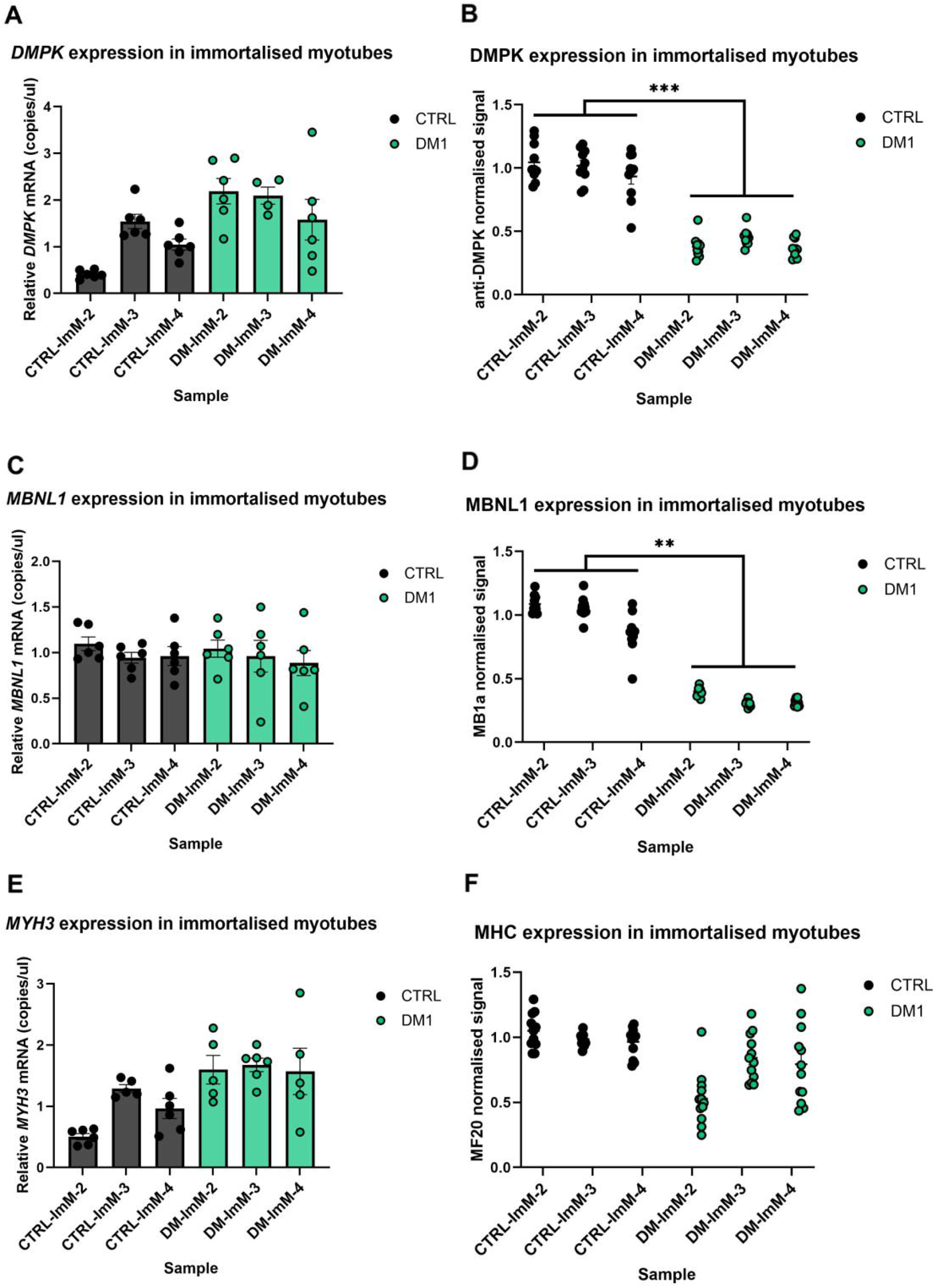
Quantification of DMPK and MBNL1 protein in control and DM1 cell cultures. *DMPK* **(A),** *MBNL1* **(C)** and *MYH3* **(E)** transcripts quantification in control and DM1-derived immortalised myotubes. Data is represented as mean ratio ± SEM between *DMPK* or *MYH3* transcripts and *HPRT1* transcripts (as loading control) or *MBNL1* transcripts and *TBP* transcripts of three technical replicates per experiment in a single experiment. DMPK **(B)**, MBNL1 **(D)** and MHC **(F)** relative protein quantification in immortalised control and DM1-derived immortalised myotubes by In-Cell Western assay. Sigma, MB1a and anti-MHC (MF20) antibody signal is normalised by Cell Tag 700 stain signal in each well. Data is represented as mean ± SEM of 3 technical replicates per experiment in three independent experiments. Significant differences were determined by two-way ANOVA and Bonferroni’s multiple comparison test (** p<0.01, ***p<0.001).

While no differences were observed at transcript level for *DMPK* and *MBNL1* (Figures 3A and 3C, respectively), DMPK and MBNL1 protein signal was lower in DM1 immortalised myotubes compared to control cultures (Figure 3B and 3D). To account for any differences in differentiation, MHC was also assessed by digital PCR and ICW, and, although differences were found, these were not significant. As described in Nuñez-Manchón et al. (2024), myoblast differentiation into myotubes is variable between different cell lines, despite being control or DM1 patient-derived cell cultures, primary or immortalised. Therefore, in this paper no statistically significant differences in MHC quantification were observed between control and patient-derived cell cultures, although variability is observed. Thischaracterisation of these cultures is consistent with the sequestration of MBNL1 protein in DMPK-expanded transcripts and the splicing alterations previously described by (30).

### DMPK and MBNL1 quantification in control and DM1-patient derived cultures

Several cell models have been used to understand the pathogenic mechanisms of DM1 and to test potential therapeutic compounds, with immortalised myotubes (28, 30, 31) and primary fibroblasts (32–34) being the most commonly used cell models. We aimed to assess these models, to evaluate their adequacy to evaluate new therapies.

In addition to the six immortalised cultures characterised in the previous figure (Figure 3), we extended our study to three DM1 and three control primary myoblast cultures purchased from Cook MyoSite, two DM1 and two control immortalised fibroblasts and three DM1 and control-derived primary fibroblasts described in (35). The differences are summarised in Figure 4 and the individual results are included in Supplementary Figures 2 (immortalised myoblasts), 3 (primary myotubes), 4 (immortalised fibroblasts) and 5 (primary fibroblasts).

**Figure 4.**
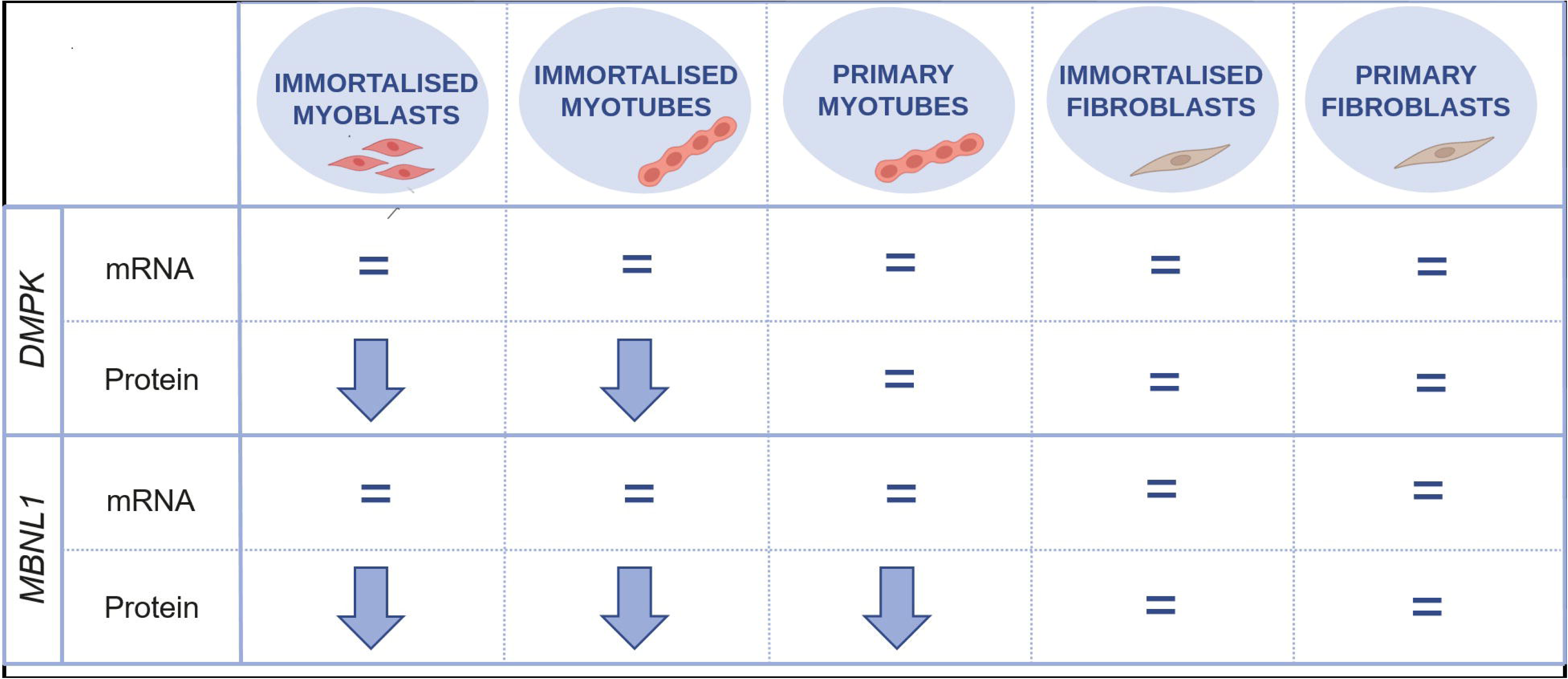
Summary of the differences in quantification of DMPK and MBNL1 between DM1 and controls, in all the cultures characterised in this project.. While no statistical differences were found at RNA levels between DM1 and controls, protein expression in myogenic lines is consistently downregulated in the DM1 patients, and ICW can pick up early downregulation of MBNL1 in primary myotubes.

DMPK protein signal was lower in DM1 immortalised myoblasts and myotubes compared to controls, while no difference was observed in primary myotubes nor immortalised or primary fibroblasts. On the other hand, MBNL1 protein signal was consistently lower in all DM1 myoblast and myotube cultures (primary and immortalised), whereas it was not altered in DM1 fibroblast cultures (neither primary nor immortalised). DMPK and MBNL1 mRNA transcripts were unaltered in all cases, suggesting that the observed differences are downstream from mRNA transcription.

### The importance of differentiation in the expression of DMPK and MBNL1

To further understand the differences we had seen between the cell models evaluated, we assessed *DMPK* mRNA expression in control primary fibroblasts and immortalised fibroblasts compared to immortalised myoblasts. While still undifferentiated, immortalised myoblasts show 3 to 10 times more *DMPK* mRNA transcripts compared to fibroblasts (Figure 5A) and, when these myoblast cultures were differentiated, and evaluated at different time points (Figure 5B), *DMPK* mRNA increased over time. We hypothesised that *DMPK* transcripts are physiologically increased in myoblasts models compared to fibroblasts, so we evaluated control and DM1 myo-inducible fibroblasts with and without MyoD induction: we observed how the induction of MyoD alone was enough to increase *DMPK* transcripts in these cultures (Figure 5C). However, DMPK protein was only downregulated in the patient’s DM1 myo-inducible cultures (Figure 5D), suggesting an increase on the expanded transcript expression and therefore, lower protein translation levels.

**Figure 5.**
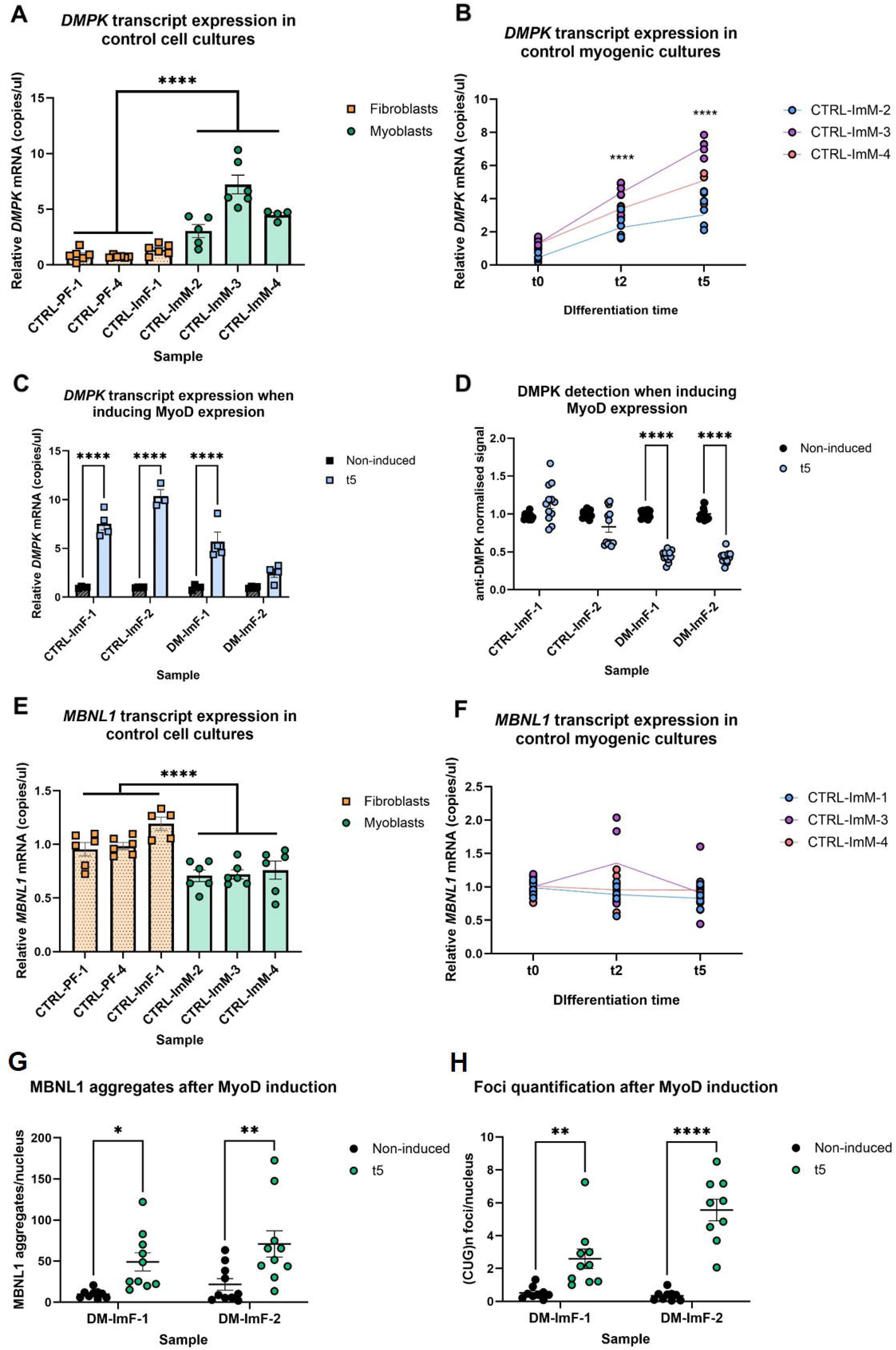
*DMPK* and *MBNL1* expression and myogenic differentiation. Expression of *DMPK* **(A and B)** and *MBNL1* **(E and F)** was evaluated by ddPCR in control fibroblasts (primary and immortalised) compared to immortalised myoblasts before (A and E) or during myogenic differentiation of the myogenic cultures (B and F). **(C)** *DMPK* mRNA and **(D)** protein expression in control myo-inducible fibroblasts with and without MyoD induction assessed by ddPCR and ICW. Transcript data is represented as mean ratio ± SEM between *DMPK* transcripts and *HPRT1* transcripts (loading control) of 2 technical replicates per experiment in three independent experiments (A and E) and 2 technical replicates per experiment in two independent experiments (C). Protein data is represented as mean ± SEM of 4 technical replicates per experiment in three independent experiments. Anti-DMPK antibody signal (800nm) is normalised by Cell Tag 700 stain signal (700nm). Significant differences were determined by two-way ANOVA and Bonferroni’s multiple comparison test (****p<0.0001). Quantification of MBNL1 aggregates (**G)** and foci **(H)** in DM1 myo-inducible fibroblasts with and without MyoD induction assessed by FISH and immunocytochemistry. Significant differences were determined by Kruskal-Wallis test followed by Dunn’s post hoc test (* p<0.05, **p<0.01, ****p<0.0001).

The increase in DMPK mRNA expression seen in myoblasts vs fibroblasts in figure 5A was accompanied by a slight but statistically significant reduced level of *MBNL1* mRNA in myoblasts compared to fibroblasts (Figure 5E). The MBNL1 mRNA expression showed no change during the myoblast maturation process (Figure 5F).

The upregulation of *DMPK* after MyoD induction suggested an increased sequestration of MBNL1 protein in expanded transcripts and, to evaluate it, we quantified MBNL1 aggregates (Figure 5G) and foci (Figure 5H) in myo-inducible DM1 fibroblasts before and after MyoD induction. Indeed, MBNL1 aggregates and foci increased after MyoD induction, suggesting that the induction increased the number of DMPK transcripts, including the expanded transcripts, and therefore caused higher MBNL1 sequestration and accumulation in foci.

### DMPK and MBNL1 quantification after treatment with DMPK-targeting antisense oligonucleotides

Most therapies targeting DM1 aim to decrease the toxicity of the *DMPK* repeats by decreasing *DMPK* transcripts (expanded and wild-type) or by avoiding the binding of MBNL1 protein to the expanded transcripts (4). However, in most cases there’s no certainty if a potentially therapeutic compound is acting on the wild-type or the expanded transcripts as, unless a specific and uncommon SNP is found in the repeated allele, it is not possible to discriminate between both alleles (36). *DMPK* expanded transcripts are thought to be retained inside the nucleus and not transferred into the cytoplasm for further translation, as opposed to WT transcripts (37). Therefore, we hypothesised that by quantifying DMPK protein, we could assess how the *DMPK* wild-type transcript is affected by DM1 potential treatments.

For proper validation of DMPK quantification, we assessed by ddPCR and ICW a siRNA targeting a *DMPK* region in exon 15 (before the 3’UTR region) (Figure 6).

**Figure 6.**
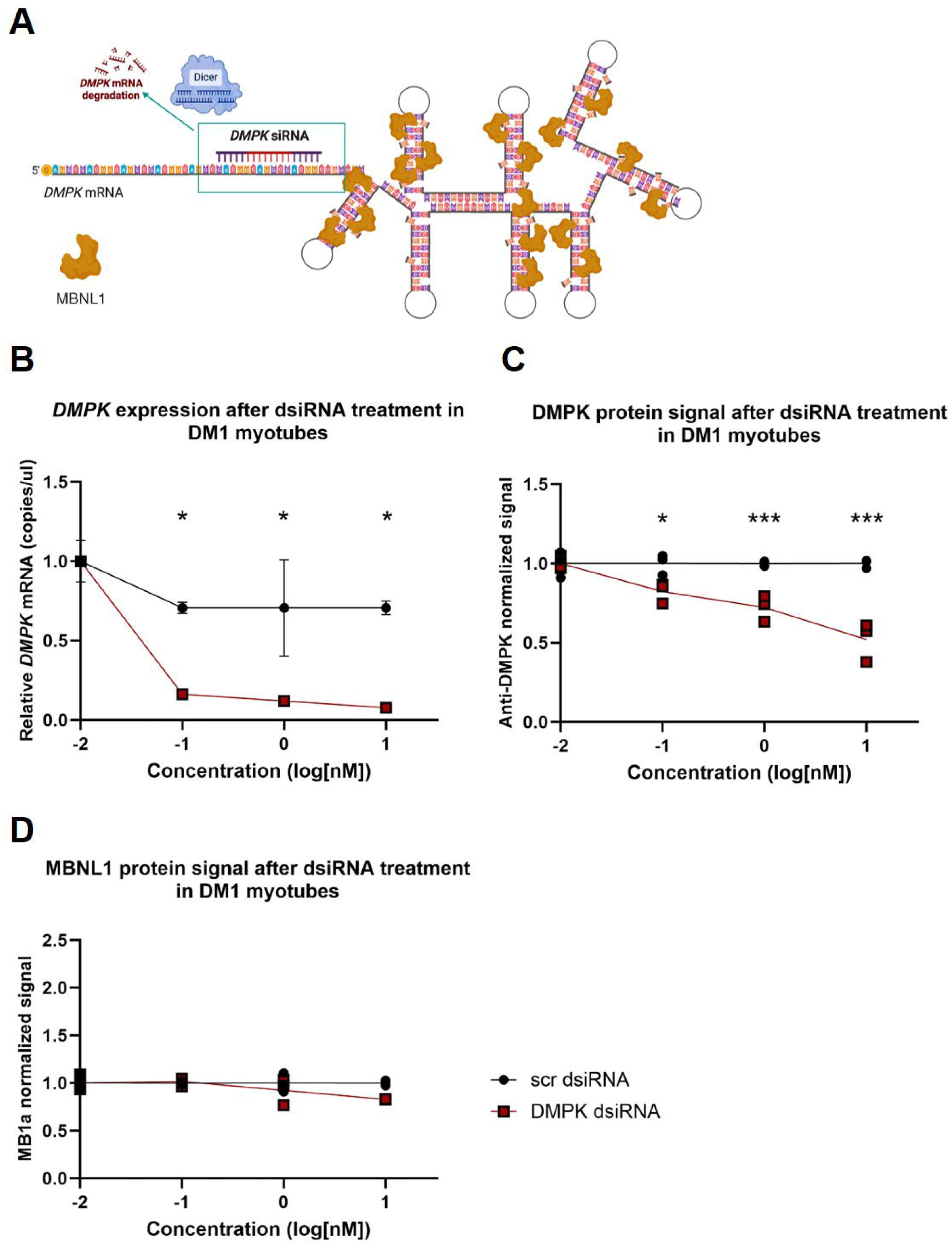
DMPK dsiRNA response quantification after differentiation. **(A)** Representation of DMPK siRNA targeting in the *DMPK* transcript. Created with Biorender.com. **(B)** Quantification of *DMPK* transcripts by ddPCR 6 days after transfection of DM1-ImM-2 myotubes with the DMPK13.1 dsiRNA and its corresponding scramble control at increasing concentrations. Data from 3 technical replicates per experiment in two independent experiments is represented as mean ratio ± SEM between *DMPK* transcripts and *HPRT1* transcripts (as loading control). Quantification by ICW of DMPK **(C)** and MBNL1 protein **(D)** 7 days after transfection of DM1-ImM-2 myotubes with the DMPK13.1 dsiRNA and scramble controls at three concentrations. Anti-DMPK and MB1a antibody signals (800nm) are normalised by Cell Tag 700 stain signal (700nm). Data from 4 technical replicates per experiment in a single experiment is represented as mean ± SEM. Significant differences were determined by two-way ANOVA and Bonferroni’s multiple comparison test (*p<0.05,** p<0.01, *** p<0.001). While DMPK expression in reduced by the dsiRNA both at RNA and protein levels, with a clear dose response at protein levels, there is no difference in the amount of MBNL1 protein quantified.

We are able to quantify a decrease in D*MPK* mRNA and protein level in DM1 immortalised myotubes after siRNA treatment, but this was not accompanied by a change in the expression of MBNL1 protein.

We also evaluated a gapmer targeting DMPK previously described in (19) (Figure 7). This *DMPK* gapmer ASO was designed to promote the degradation of *DMPK* transcripts through RNase H1 by targeting a sequence in exon 15 of *DMPK* mRNA, the 3’UTR region, nearby the region targeted by the siRNA evaluated in figure 6. In their publication (Figure 7A), 24-48h after treatment of the cultures with the DMPK gapmer at 100nM, the authors observed by qPCR a decrease in *DMPK* transcripts, by immunohistochemistry a decrease of MBNL1 density in foci, and an improvement of overall missplicing. We completed their evaluation by quantifying *DMPK* transcripts by ddPCR (Figure 7B) and DMPK and MBNL1 protein by ICW. Our quantification aligns with their results and showed a decrease in *DMPK* transcripts at all concentrations, a moderate decrease in DMPK protein signal (Figure 7C) and an increase of up to twice the amount of MBNL1 protein, compared to the scramble control (Figure 7D), possibly indicating the release of MBNL1 protein from foci.

**Figure 7.**
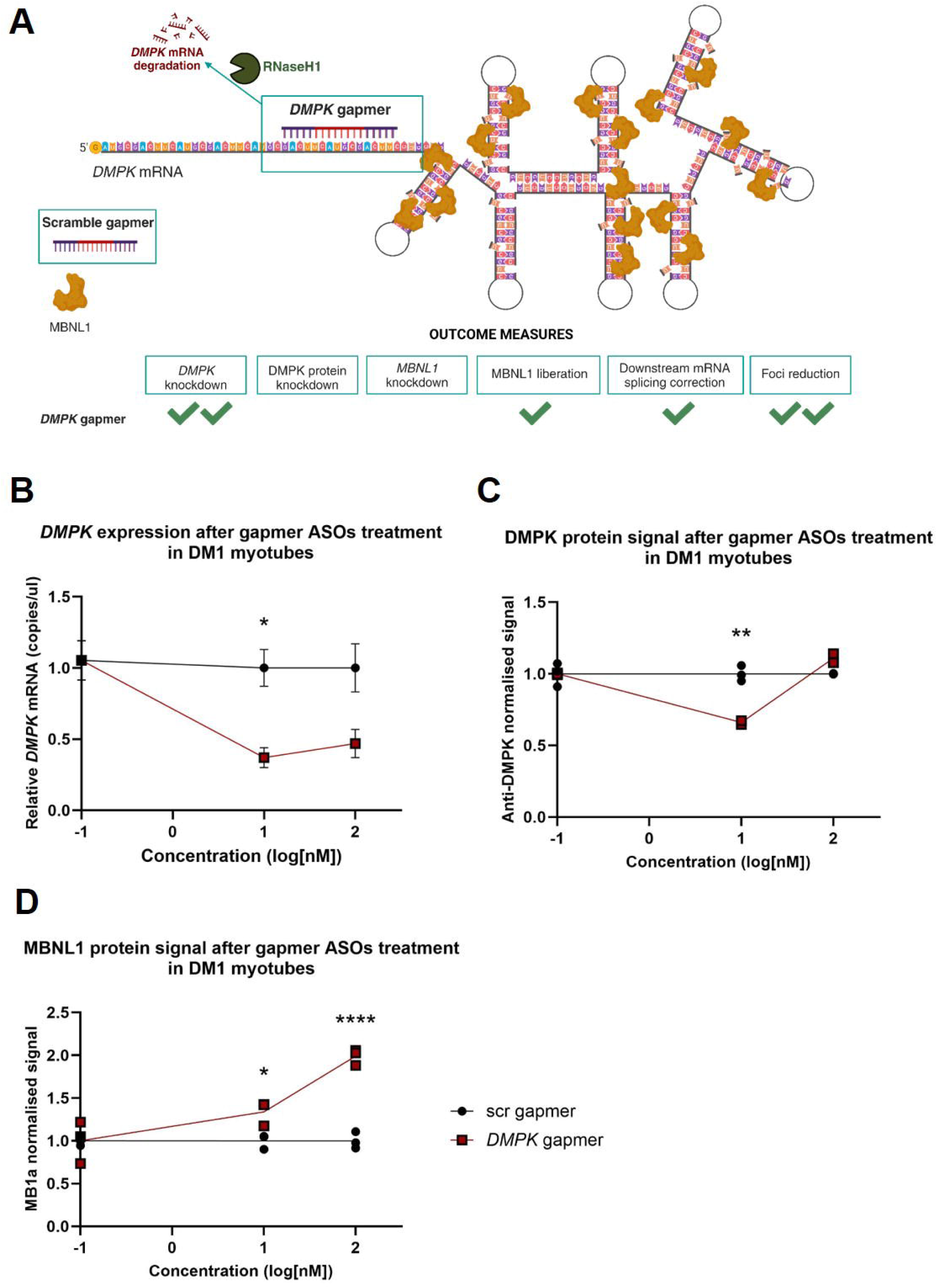
DMPK gapmer effect on DM1 myoblasts and myotubes. **(A)** Representation of DMPK gapmer targeting in the *DMPK* transcript and summary of the response to treatment with 100nM of this gapmer of DM-ImM-1 myoblasts as previously reported by (El Boujnouni et al., 2023). Created with Biorender.com. **(B)** Quantification by ddPCR of *DMPK* transcripts 6 days after transfection of DM1-ImM-1 myotubes with the DMPK gapmer and its corresponding scramble control at increasing concentrations. Data of three technical replicates per experiment in a single experiment is represented as mean ratio ± SEM between DMPK transcripts and HPRT1 transcripts (as loading control). **(C)** Quantification by ICW of DMPK and MBNL1 protein **(D)** 7 days after transfection of DM1-ImM-2 myotubes with the DMPK gapmer and scramble control at different concentrations. Anti-DMPK and MB1a antibody signals (800nm) are normalised by Cell Tag 700 stain signal (700nm). Data of 3 technical replicates per experiment in two independent experiments is represented as mean ± SEM. Significant differences were determined by two-way ANOVA and Bonferroni’s multiple comparison test (*p<0.05,** p<0.01,****p<0.0001). The DMPK gapmer was succesful at both reducing *DMPK* expression and increasing the amount of MBLN1 protein detected, in a dose dependent manner.

## Discussion

Myotonic dystrophy type 1 (DM1) is a multisystemic disease with molecular dysfunctions affecting various tissues. Research is hindered by the limited availability of patient cell lines: fibroblasts, more easily obtained than myoblasts, are preferred for their accessibility and robust culture conditions, and, although they might reproduce some of the splicing defects that are hallmarks of the disease, other hallmarks may not be easily found in this model. Foci, containing RNA repeats and MBNL1 protein, are a key element of DM1 pathology and are widely used as an outcome measure by many research groups. Therefore, proper detection of RNA and MBNL1 foci is critical in DM1 research. Many therapies in DM1 research target *DMPK* transcripts and aim reducing MBNL1 binding and therefore, reducing MBNL1 sequestration by either reducing the expanded *DMPK* transcripts or displacing MBNL1 from them. Being able to quantify both the DMPK transcripts and free MBNL1, that is, the MBNL1 protein that it is not trapped in the expanded transcripts, particularly in a high-throughput format is a useful tool to deepen in the biology of the disease, characterise the models used in the field and screen for potential treatments.

We have developed a platform for quantifying DMPK and MBNL1 expression in various cell models, with the goal of identifying biomarkers for *in vitro* drug screening and enhancing therapeutic development for DM1. With this methodology, we have characterised a large cohort of cell models widely used in the field, including immortalised and primary myoblasts, myotubes, and fibroblasts, to explore how DM1 may alter *DMPK* and *MBNL1* expression in these cultures. DMPK protein levels were reduced in immortalised myoblasts and myotubes but remained unchanged in primary myotubes and fibroblasts. The lack of DMPK depletion in primary myotubes might be due to the longer expansions in the DM1 immortalised cultures vs the primary ones. It is likely that transcript sequestration is more prominent in samples where there are either longer expansions or a larger amount of these mRNAs, as seen in myotubes, where longer repeat lengths might exacerbate transcript sequestration (38). Furthermore, *DMPK* mRNA levels were higher in immortalised myoblasts compared to fibroblasts, and these transcripts increased during myogenic differentiation. Additionally, in myo-inducible models, protein levels did not correlate with the increase in *DMPK* transcripts, especially in DM1 cells, where DMPK protein levels were reduced, suggesting higher transcript sequestration.

Similarly, we also observed a reduction in MBNL1 protein levels in DM1 myoblasts and myotubes compared to controls, consistent with previous reports. No difference was observed between DM1 and control fibroblasts. Interestingly, MBNL1 protein levels were lower in myoblasts compared to fibroblasts and no changes were observed during myogenic differentiation. These findings suggested that DM1 myoblasts and myotubes may experience greater alterations due to the combined effects of higher *DMPK* transcript levels and reduced MBNL1 availability. Further analysis in myo-inducible models showed increased MBNL1 aggregates and more RNA foci in DM1 myotubes compared to fibroblasts, indicating higher accumulation of *DMPK*-expanded transcripts in muscle cells.

Our platform enabled the assessment of *DMPK* and MBNL1 as potential outcome measures for drug testing in DM1. The results indicated that myoblasts and myotubes were more suitable than fibroblasts for screening molecules targeting *DMPK* and MBNL1. Moreover, the quantification of *DMPK* mRNA and protein could distinguish between compounds that affect expanded versus non-expanded DMPK transcripts, providing insight into therapeutic mechanisms. Treatment with *DMPK*-targeting molecules, such as the gapmer and dsiRNA, showed differential effects: the DMPK gapmer decreased *DMPK* transcripts without reducing protein levels but increased MBNL1 protein, while the dsiRNA reduced both *DMPK* transcripts and protein, with no effect on MBNL1. These differences highlight the distinct mechanisms of action of both ASOs, with the gapmer acting at the pre-mRNA level to release MBNL1 from sequestration and the dsiRNA acting at post-transcriptional mRNA level in the cytoplasm (39). RISC and DICER are located in the cytoplasm of human cells and, although with different delivery methods siRNAs are able to decrease *DMPK* transcripts, in our hands we observed how this *DMPK* silencing is targeting the *DMPK* wild-type transcript. Then, reducing DMPK protein, and not showing any effect at MBNL1 liberation, as it is not able to target the *DMPK* expanded transcript.

In conclusion, in this manuscript, we describe an effective platform for assessing DMPK and MBNL1 in cell culture, we characterise a large cohort of DM1 cell culture models widely used in preclinical research and corroborate the screening capabilities of our platform by confirming and completing the effect of previously reported potential therapies. We expect our results to be of use to the DM1 research community.

## Materials and methods

### Myoblasts cultures

All cell cultures are detailed in Table 1. Immortalised myoblasts were immortalised by the MyoLine platform (Paris) and kindly provided by Dr. Furling (Institut of Myology) and Dr. Nogales-Gadea from Germans Trias I Pujol Research Institute (IGTP). DM-ImM-1 culture was previously characterised (40) and (CTG)n repeats were sized by Southern blot (see Table 1). DM-ImM-2, DM-ImM-3 and DM-ImM-4 cultures were characterised in (30). (CTG)n repeats were evaluated by small-pool PCR followed by Southern blot. In this paper, DM-ImM-2 corresponds to JCC-DM1, DM-ImM-3 to GPM-DM1 and DM-ImM-4 to ADE-DM1 in (30). Primary myoblasts were purchased from Cook MyoSite (USA) which also provided with the range of (CTG)n repeats, quantified in the patient’s blood samples by Southern blot.

**Table 1.**
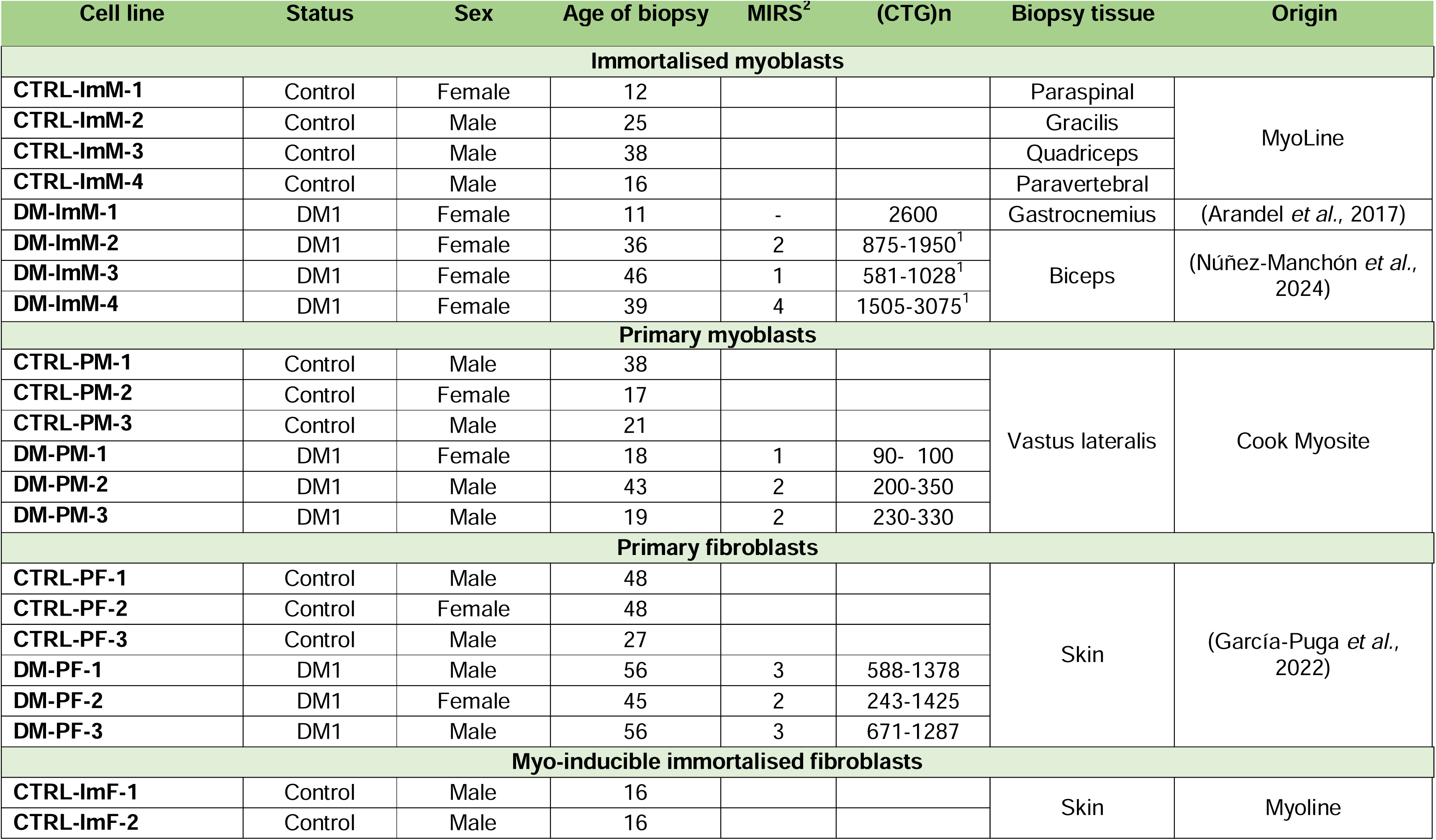
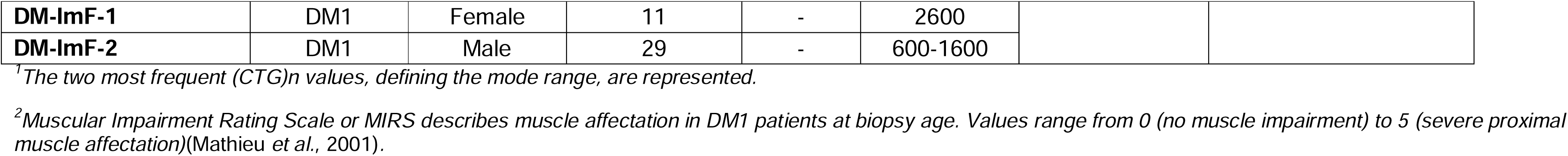
Myoblasts and fibroblasts cultures used in this publication.

All myoblast cell lines were mantained in growth medium, consisting of 1:1 Skeletal Muscle Cell Growth Medium (SMM, PELOBiotech, Germany) and Dulbecco’s Modified Eagle medium (DMEM, Corning®, USA) supplemented with 10% foetal bovine serum (FBS), 2% GlutaMax and 1% Penicillin/Streptomycin (PenStrep). All supplements were purchased from Gibco™ (USA). Cell cultures were only used up to passage 10, to avoid changes in expansion size due to cell passage. Myoblast cultures were incubated at 37°C in 5% CO_2_ and medium was refreshed every 48-72 hours. For myogenic differentiation, myoblasts were seeded in growth medium in plates coated with 1% Matrigel (Corning®) until they reached 80-90% confluence, usually after 24-48 hours. Medium was then replaced with differentiation medium (DM). DM was prepared with high-glucose DMEM containing GlutaMax (Gibco™), 1% PenStrep, 1% KnockOut™ Serum Replacement (Gibco™) and 1% insulin-transferrin-selenium-ethanolamine (ITS-X from Gibco™). Myoblasts were cultured in DM until myotubes were observed: typically, they were harvested on day 6 for mRNA studies, and on day 7 for protein studies: pelleted for quantification by Jess simple western and fixed for In-Cell Western, FISH and immunofluorescence.

### Fibroblasts cultures

Primary fibroblasts (see Table 1) were kindly provided by Dr. Lopez de Munain (BioGipuzkoa HRI). MIRS was evaluated at age of biopsy and CTG expansions were sized by small-pool PCR and Southern blot (35). Myo-inducible immortalised fibroblasts had been generated by the MyoLine platform (Paris) and were kindly provided by Dr. Furling (Institut of Myology). DM-ImF-1 and DM-ImM-1 were derived from biopsies from the same patient. DM-ImF-2 repeat size was measured by small-pool PCR and Southern blot.

Primary and immortalised fibroblast cultures were maintained in DMEM (Corning®) supplemented with 10% FBS (Gibco™), 1% PenStrep (Gibco™) and 2% GlutaMax (Gibco™). They were incubated at 37°C in 5% CO_2_ and the medium was refreshed every 48-72 hours. For MyoD induction of myo-inducible immortalised fibroblasts, cells were maintained in fibroblast medium until cell passage, then, they were resuspended in DM containing 0.02% of doxycycline (Merck, USA) and cultured at 37°C and 5% CO_2_ for 5 days. The medium was refreshed every 48 hours, and doxycycline was added to the medium just before it was added to the cells.

### ASOs treatment

Two different kinds of ASOs were used, dsiRNAs and gapmers. All were purchased lyophilised from Integrated DNA Technologies (IDT, USA). DMPK13.1 dsiRNA and its scramble control were resuspended in duplex-buffer as per manufacturer’s instructions, while gapmers ASOs were resuspended in MiliQ water in sterile conditions. Sequences are detailed in Table 2.

**Table 2.**
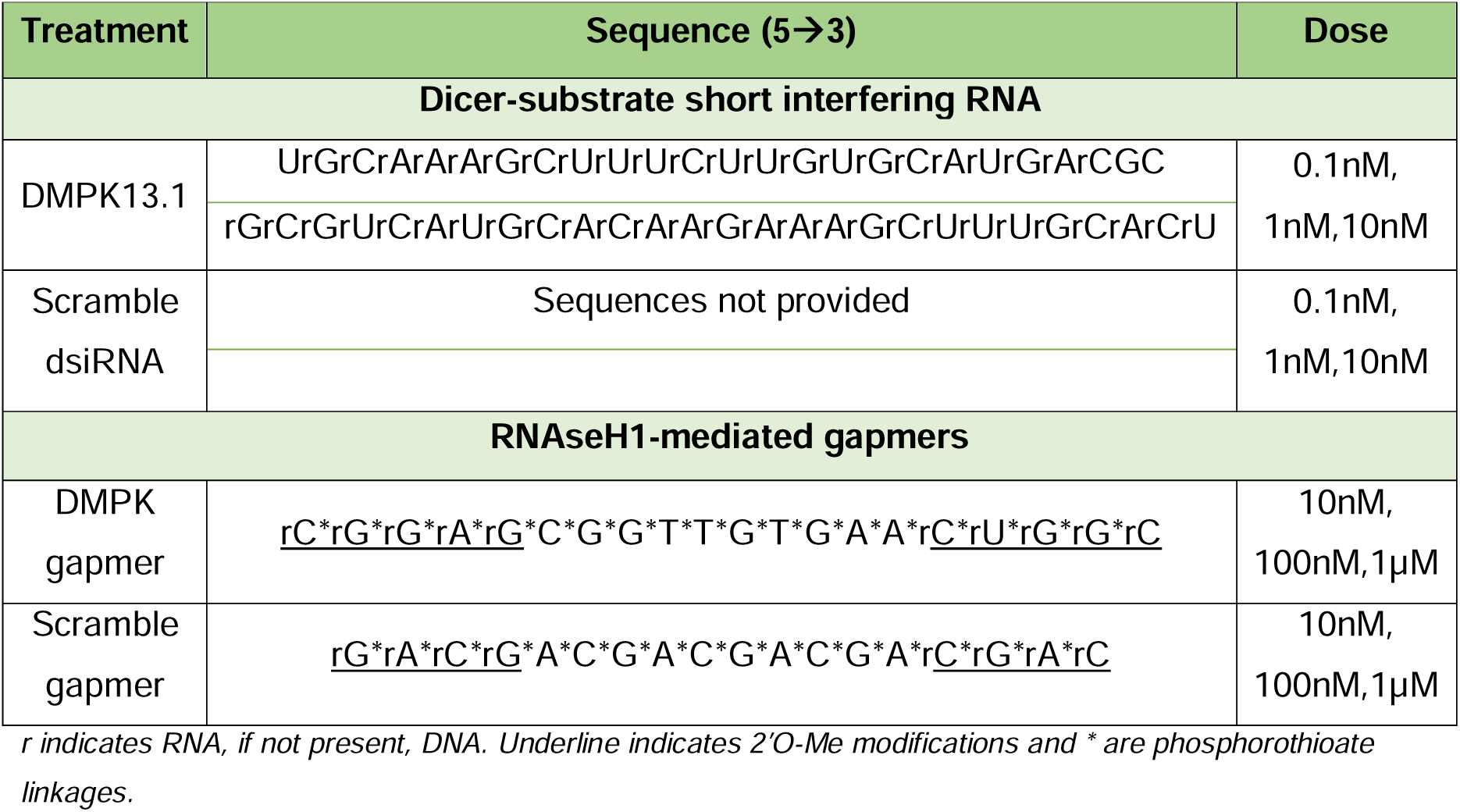
Antisense oligonucleotides used in this publication.

All ASOs were transfected into myoblasts using Lipofectamine 3000 (Invitrogen™, USA) 24h after seeding, at 70-80% confluence, following the manufacturer’s protocol. Fresh growth medium was added 4h after transfection and was replaced with DM when cultures reached 80-90% confluence, typically 24-48 hours after transfection. Myoblasts were then incubated at 37°C in 5% CO_2_ for 6 or 7 days until myotubes were visually observed. Medium was refreshed with fresh DM every 2-3 days until harvest or fixation.

### Fluorescent in situ hybridisation (FISH)

FISH is the standard method used to visualise foci in cultures (41). Briefly, cells were fixed with 10% formalin solution (Sigma) for 10 minutes followed by three washes of 5 minutes with PBS. Then, cells were pre-hybridised with 30% formamide in 2xSSC for 10 minutes at RT. Meanwhile, an hybridisation buffer was prepared containing 40% formamide, 10% saline sodium citrate buffer (SSC) 20X, 20% bovine serum albumin (BSA) 1%, 0.1g/ml dextran sulphate, 10% vanadyl complex 20mM, 10% yeast tRNA (10ug/ml), 10% herring sperm DNA and a (CAG)_7_-Cy3 probe (1/100) purchased from IDT. The samples were hybridised overnight at 37°C. The following day, samples were washed twice with 2x SSC in 30% formamide for 15 min at 42°C in an incubation oven. The hybridised samples were then used for immunocytochemistry.

### Immunocytochemistry

All samples (hybridised samples from FISH or fixed samples for direct immunofluorescence) were permeabilised with 0.1% Triton X-100 in PBS and blocked with Intercept® (PBS) Blocking Buffer (LI-COR) for 2 hours. After blocking samples were incubated overnight with primary antibodies at 4°C at the appropriate dilutions: MB1a (The MDA Monoclonal Antibody Data Source), anti-desmin (Abcam) or anti-DMPK (HPA007164 from Sigma). The following day, coverslips were washed with 0.1% Tween in PBS and incubated with a secondary antibody mixture (Alexa Fluor™ 647 goat anti-Rabbit IgG (H+L) Secondary Antibody and Alexa Fluor™ 488 goat anti-Mouse IgG (H+L) Secondary Antibody). All secondary antibodies were purchased from Life Technologies. Coverslips were washed again with 0.1% Tween in PBS and incubated for 10 minutes with Hoescht 3342 (at 1/1000) from Life Technologies. Coverslips were then mounted with ProLong™ Diamond Antifade Mountant. Images were captured using a Zeiss LSM 880 ID SCAN microscope at 40x for DMPK antibody screening and 63x for foci and MBNL1 quantification and analysed using ZEN black software and Fiji.

### In-Cell western

For fibroblasts analysis, 4000 fibroblasts/well were seeded into a well of 96-well plates. Cells were cultured for 72 hours prior to fixation. For myoblasts and myotubes, 7500 cells/well were seeded into a well of 96-well plate. Myoblasts were cultured for 48 hours prior to fixation for myoblast protein detection. For myotube differentiation, myoblasts were cultured for 24-48 hours before the addition of differentiation medium until they reached 80-90% confluence. Myotubes were then fixed on day 7 of differentiation. Myo-inducible fibroblasts were cultured for 5 days prior to fixation, with the differentiation medium replaced with fresh doxycycline every 48 hours. Cells were fixed with 10% formalin solution for 10 minutes and 3 washes of 5 minutes with PBS 1X. After fixation, plates were permeabilised with 0.1% Triton X-100 in PBS and blocked for 2 hours with Intercept® (PBS) Blocking Buffer (LI-COR). Primary antibodies were incubated overnight at 4°C at the respective dilutions (see *Immunocytochemistry*). The following day, plates were washed with 0.1% Tween in PBS and incubated with the corresponding secondary antibody mixtures consisting in Cell Tag 700 Stain and either IRDye® 800CW Goat anti-Mouse IgG Secondary Antibody or IRDye® 800CW Goat anti-Mouse IgG Secondary Antibody. Secondary antibody dilutions were described in the manufacturer’s protocol. All reagents were supplied by LI-COR Biosciences. Plates were washed again with 0.1% Tween in PBS and 200ul/well of PBS were added before scanning the plates in an Odyssey® M Imager (LI-COR Biosciences). Data was then obtained with LI-COR Acquisition Software and analysed with Empiria Studio® Software before statistical analysis.

### Jess simple western

Protein was extracted from myotube cultures following RIPA extraction and lysis buffer (Thermo Scientific™, USA) supplemented with protease and phosphatase inhibitor cocktails (Roche, Switzerland) following manufacturer’s protocol and later quantified using a Pierce™ BCA Protein Assay Kit (Thermo Scientific™) following the manufacturer’s protocol. Quantification was performed in an Infinite® 200 PRO plate reader (TECAN) at 562nm. For immunodetection, 3ul of 0,4mg/ml sample was loaded into a 12-220 kDa Separation Module (Bio-Techne, France), together with the appropriate antibodies and blocking buffer. All reagents were purchased from Bio-Techne. The cartridge was then read on a Jess instrument and the data was analysed using Bio-Techne’s Compass SW software.

### Digital droplet PCR

RNA was extracted from cell pellets using the RNeasy Micro Kit for pellets collected from 12-well plates and the RNeasy Mini Kit for pellets collected from 6-well plates, following manufacturer’s protocols (QIAGEN, Germany). Once extracted, RNA was quantified using a Nanodrop apparatus and 100ng-1µg of the extracted RNA was used as template for reverse transcription with SuperScript™ IV Reverse Transcriptase (Invitrogen™), following manufacturer’s guidelines. The resulting cDNA was diluted to 0,05ng/µl in nuclease-free water for the ddPCR protocol. 2 µl of cDNA were used in 20µl PCR reactions, that included 10µl of ddPCR™ Supermix for Probes (No dUTP) from Bio-Rad, 7µl of nuclease free-water, 0.5µl of PrimeTime™ qPCR Probe Assay (IDT) marked with FAM, 0.5µl of PrimeTime™ qPCR Probe Assay (IDT) marked with HEX. Each reaction included a PrimeTime™ qPCR Probe Assay (IDT) targeting *HPRT1* or *TBP* as loading control. Housekeeping genes in each case were selected based on expression levels. To generate droplets, 20µl of the prepared ddPCR reactions and 70µl of the Droplet Generation Oil for Probes (Bio-Rad) were added to the 8-channel droplet generator cartridge (Bio-Rad) sealed with a DG8 gasket (Bio-Rad) and placed in the QX200 droplet generator (Bio-Rad). 40µl of the droplet mixture were collected from the cartridge and transferred to a 96-well semi-skirted ddPCR plate (Bio-Rad), sealed with a pierceable foil and amplified on a deep-well thermal cycler using the following conditions: enzyme activation for 10 min at 94°C followed by 40 cycles of denaturation for 30 seconds at 94°C, 1 min at 59°C for annealing and extension, and heat deactivation for 10 min at 98°C. Plates containing the amplified droplets were then analysed in the QX200 droplet reader. Results were analysed using QuantaSoft™ Software and QuantaSoft™ Analysis Pro (Bio-Rad) software. Droplet number was used as a quality control and wells with more than 10.000 droplets were included in the analysis.

### Statistical analysis

All results were expressed as mean ± standard error of the mean (SEM). Statistical analysis was performed using GraphPad Prism 10.1.2 software. Data distribution was assessed using the Shapiro-Wilk test. If the normality test was passed, outliers were detected using the ROUT method (Q=1%) and heteroscedasticity was assessed using the Fisher test. Normal data with equal variance was assessed by parametric method, one-way analysis of variance (ANOVA) followed by Bonferroni’s multiple correction test. Data that did not follow normal distribution or equality of variances was analysed by non-parametric tests, Kruskal-Wallis test followed by Dunn’s post hoc test. P-values used in this study to determine statistical significance where as follows: *p-value <0.05, **p-value<0.01, ***p-value<0.001, ****p-value<0.0001.

## Acknowledgements

We acknowledge the patients that donated skin samples origin of the cell cultures provided by Dr López de Munain, BioGipuzkoa-BioDonostia Health Research Institute (Donostia-San Sebastián, Spain), and the Institut de Myologie (Paris, France). We gratefully acknowledge the MB1a antibody provided by Professor Glenn Morris from the Muscular Dystrophy Association (MDA) Monoclonal Antibody Resource, which distributes antibodies for research in neuromuscular diseases worldwide from Oswestry, United Kingdom. We would also like to thank Dr. Najoua El Boujnouni and Dr. Derick G. Wansink, Radboud university medical center (Nijmegen, the Netherlands), for the opportunity of testing their oligonucleotides while a research stay in their laboratory, and Dr. Pešović, Centre for Human Molecular Genetics of the University of Belgrade (Serbia) for the sizing of DM-ImF-2 repeats.

## Data and resource availability

All relevant data can be found within the article and its supplementary information. All materials and further information of this study is available upon request.

## Figure and table legends

**Table 3. Myoblasts and fibroblasts cultures used in this publication.**

1The two most frequent (CTG)n values, defining the mode range, are represented.

2Muscular Impairment Rating Scale or MIRS describes muscle affectation in DM1 patients at biopsy age. Values range from 0 (no muscle impairment) to 5 (severe proximal muscle affectation)(Mathieu et al., 2001).

**Table 4. Antisense oligonucleotides used in this publication.**

r indicates RNA, if not present, DNA. Underline indicates 2’O-Me modifications and * are phosphorothioate linkages.

**Supplementary figure 8. Optimisation of In-Cell western. (A)** Different cell densities of CTRL-ImM-1 were seeded in a 96-well plate in triplicates (2000cells/well, 3500 cells/well, 5000 cells/well, 7500 cells/well, 9000 cells/well and 10500 cells/well) and incubated for 48h. Then, the plate was fixed with formalin solution and stained with Cell Tag 700 Stain. Signal intensity after staining CTRL-ImM-1 cell cultures with different antibody dilutions of **(B)** Sigma’s anti-DMPK antibody and **(C)** MB1a antibody was measured by In-Cell western. Data is represented as mean ± SEM of 4 technical replicates per experiment in a single experiment. Analysis was performed with Empiria Studio Software. DMPK **(D)** and MBNL1 **(E)** protein quantification 48h after seeding. Sigma and MB1a antibody signal (800nm) is normalised by Cell Tag 700 stain signal (700nm). Protein data is represented as mean ± SEM of 4 technical replicates per experiment in three independent experiments. Significant differences were determined by two-way ANOVA and Bonferroni’s multiple comparison test (***p<0.001).

**Supplementary figure 9. DMPK and MBNL1 quantification in immortalised control and DM1 myoblasts.** DMPK **(A)** and MBNL1 **(C)** transcripts quantification by ddPCR 24h after seeding. Data is represented as ratio between DMPK transcripts and HPRT1 transcripts as loading control or MBNL1 transcripts and TBP transcripts as loading control. Transcripts’ data is represented as mean ± SEM of 2 technical replicates per experiment in three independent experiments. DMPK **(B)** and MBNL1 **(D)** protein quantification 48h after seeding. Sigma and MB1a antibody signal (800nm) is normalised by Cell Tag 700 stain signal (700nm). Protein data is represented as mean ± SEM of 4 technical replicates per experiment in three independent experiments. Significant differences were determined by two-way ANOVA and Bonferroni’s multiple comparison test (***p<0.001).

**Supplementary figure 10. MHC, DMPK and MBNL1 quantification in primary control and DM1 myotubes.** MYH3 **(A)**, DMPK **(C)** and MBNL1 **(E)** transcripts quantification by ddPCR seven days after differentiation. Data is represented as ratio between MYH3 or DMPK transcripts and HPRT1 transcripts as loading control or MBNL1 transcripts and TBP transcripts as loading control. Transcripts’ data is represented as mean ± SEM of 2 technical replicates per experiment in three independent experiments. MHC **(B)**, DMPK **(D)** and MBNL1 **(F)** protein quantification 48h after seeding. MF20, Sigma and MB1a antibody signal (800nm) is normalised by Cell Tag 700 stain signal (700nm). Protein data is represented as mean ± SEM of 4 technical replicates per experiment in three independent experiments. Significant differences were determined by two-way ANOVA and Bonferroni’s multiple comparison test (***p<0.001).

**Supplementary figure 11. DMPK and MBNL1 quantification in immortalised control and DM1 fibroblasts.** DMPK **(A)** and MBNL1 **(C)** transcripts quantification by ddPCR 24h after seeding. Data is represented as ratio between DMPK transcripts and HPRT1 transcripts as loading control or MBNL1 transcripts and TBP transcripts as loading control. Transcripts’ data is represented as mean ± SEM of 2 technical replicates per experiment in three independent experiments. DMPK **(B)** and MBNL1 **(D)** protein quantification 48h after seeding. Sigma and MB1a antibody signal (800nm) is normalised by Cell Tag 700 stain signal (700nm). Protein data is represented as mean ± SEM of 4 technical replicates per experiment in three independent experiments. Significant differences were determined by two-way ANOVA and Bonferroni’s multiple comparison test (***p<0.001).

**Supplementary figure 12. DMPK and MBNL1 quantification in primary control and DM1 fibroblasts.** DMPK **(A)** and MBNL1 **(C)** transcripts quantification by ddPCR 24h after seeding. Data is represented as ratio between DMPK transcripts and HPRT1 transcripts as loading control or MBNL1 transcripts and TBP transcripts as loading control. Transcripts’ data is represented as mean ± SEM of 2 technical replicates per experiment in three independent experiments. DMPK **(B)** and MBNL1 **(D)** protein quantification 48h after seeding. Sigma and MB1a antibody signal (800nm) is normalised by Cell Tag 700 stain signal (700nm). Protein data is represented as mean ± SEM of 4 technical replicates per experiment in three independent experiments. Significant differences were determined by two-way ANOVA and Bonferroni’s multiple comparison test (***p<0.001).

