## Supplementary figures and images for "Characterisation of *DMPK* and MBNL1 expression in cell models of Myotonic Dystrophy: A platform for drug screening"

### Supplementary figure 1

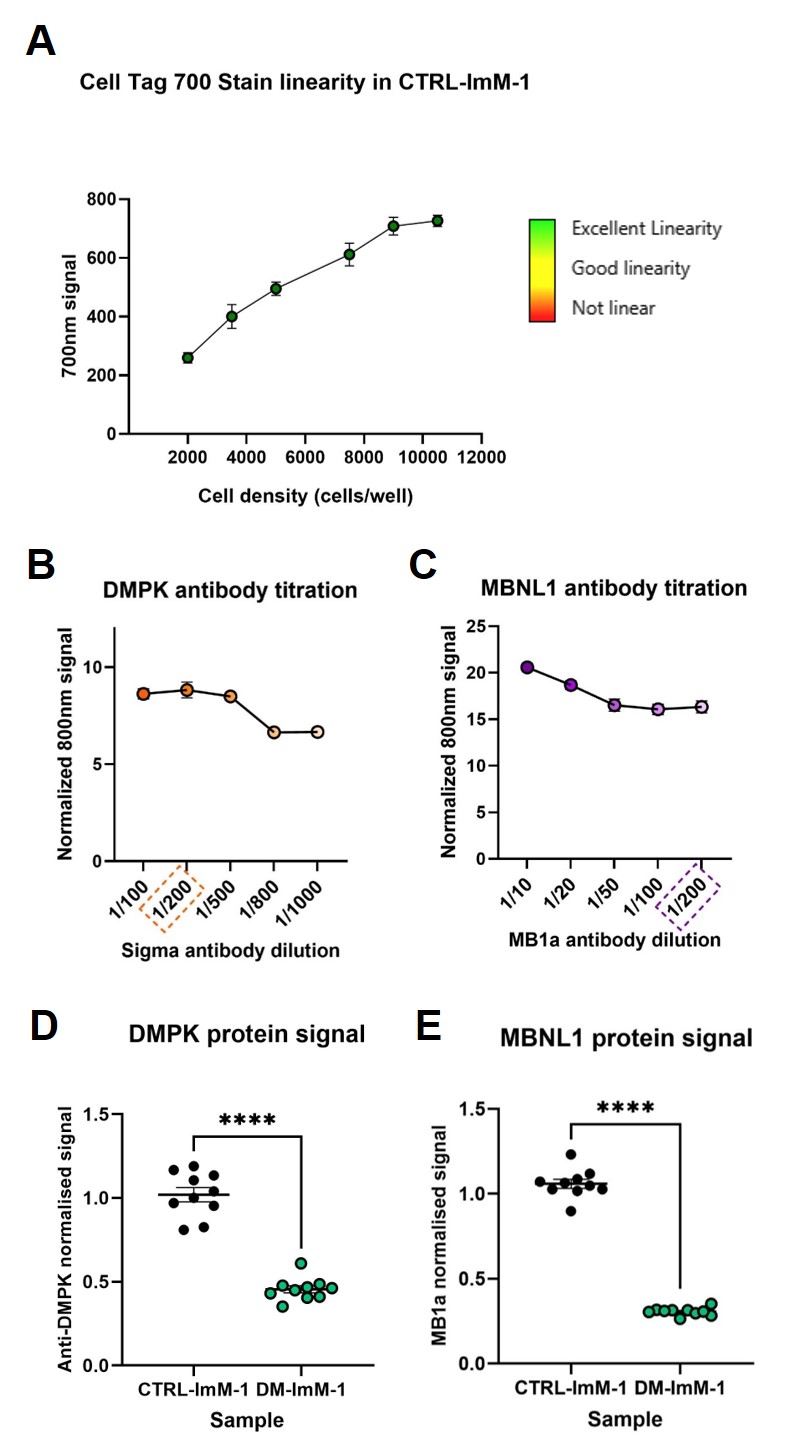

### Supplementary figure 2

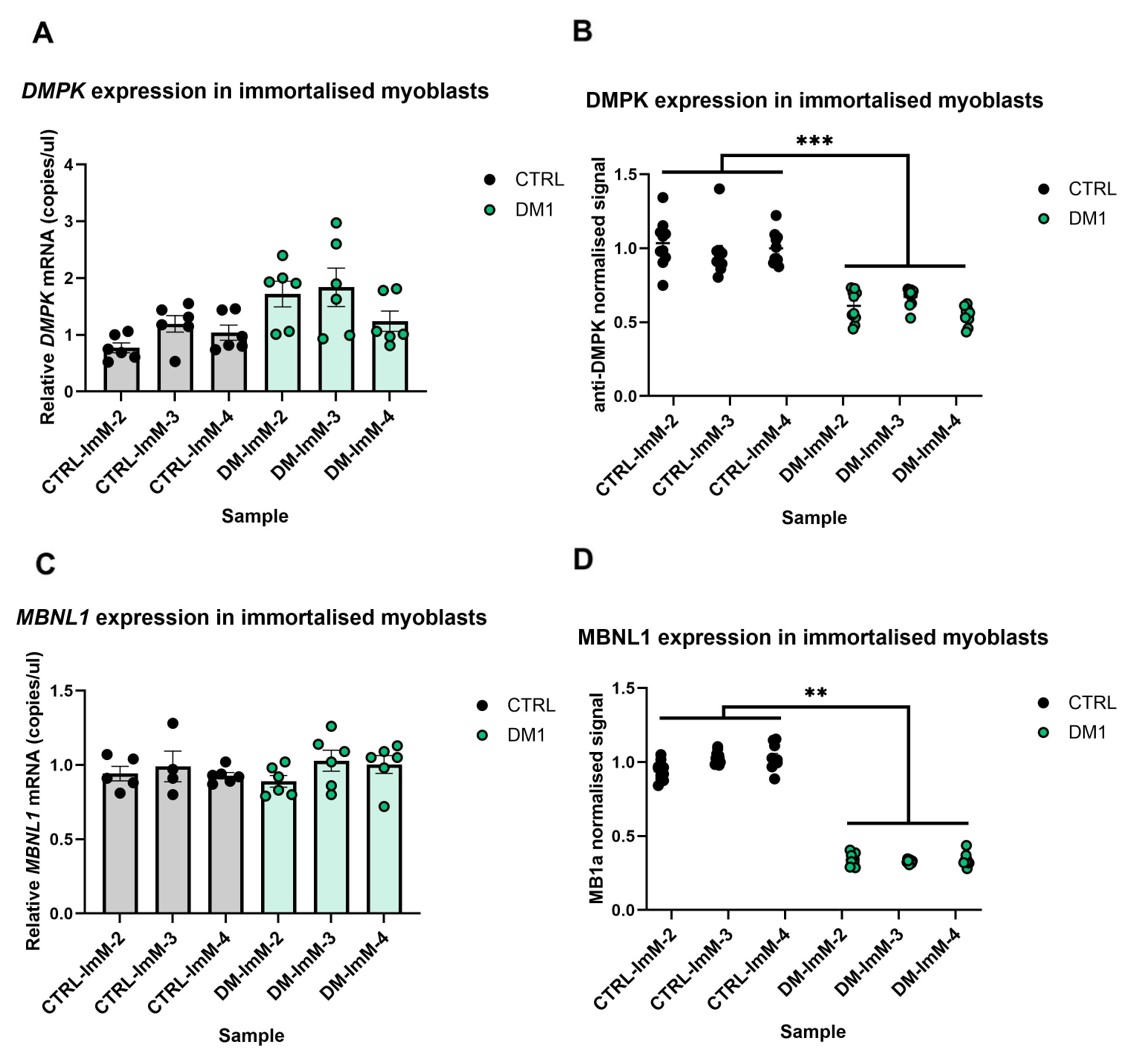

### Supplementary figure 3

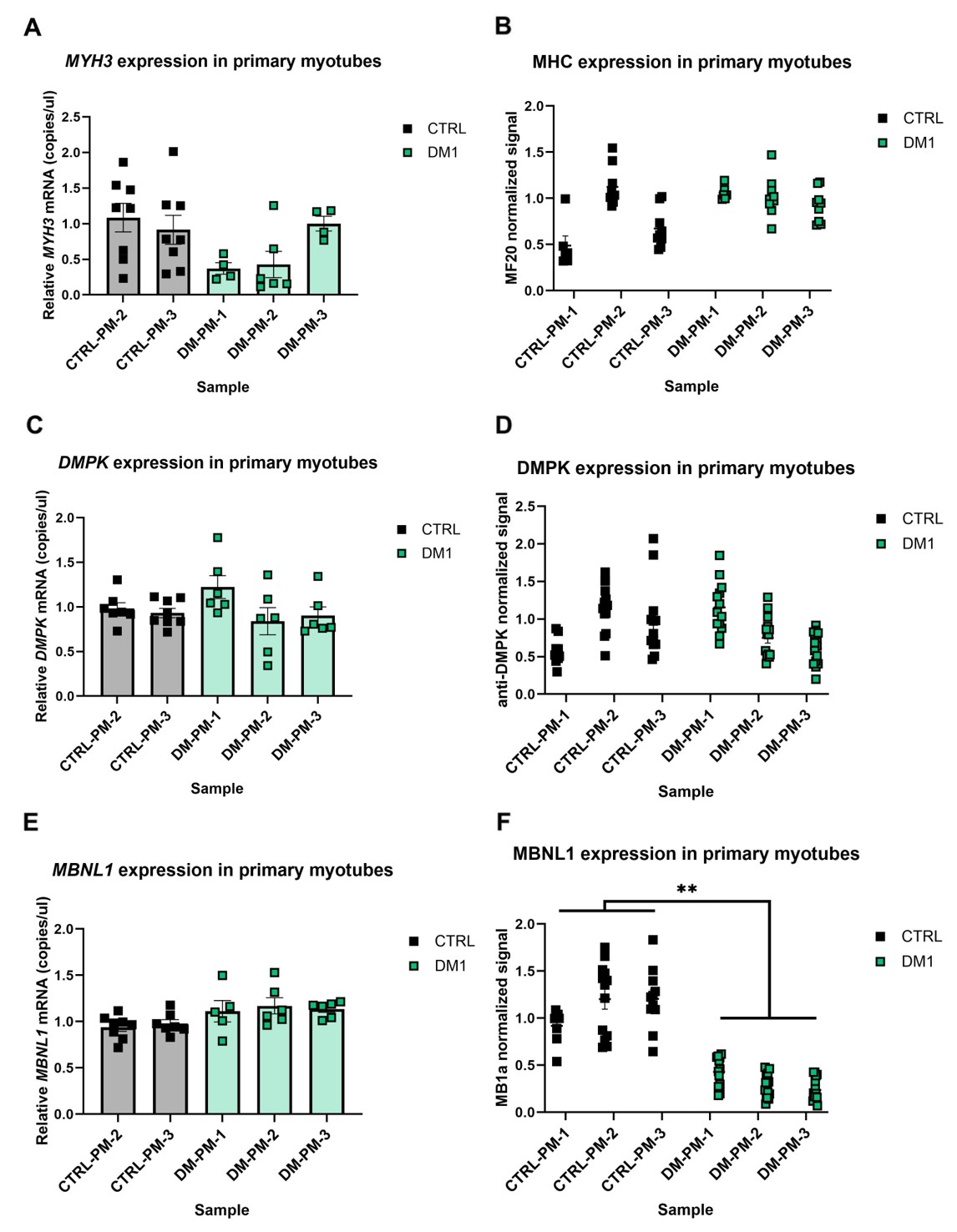

### Supplementary figure 4

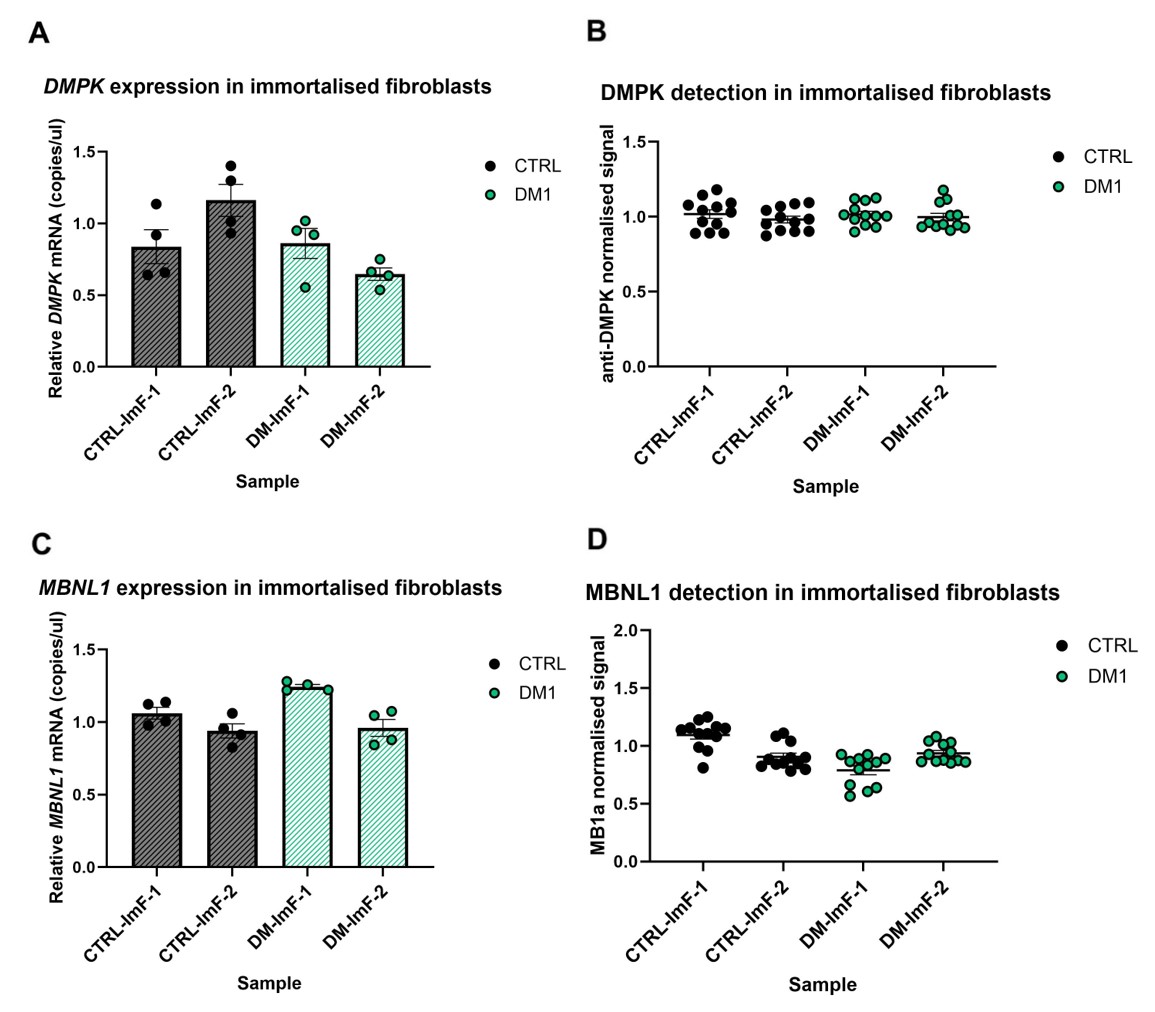

### Supplementary figure 5

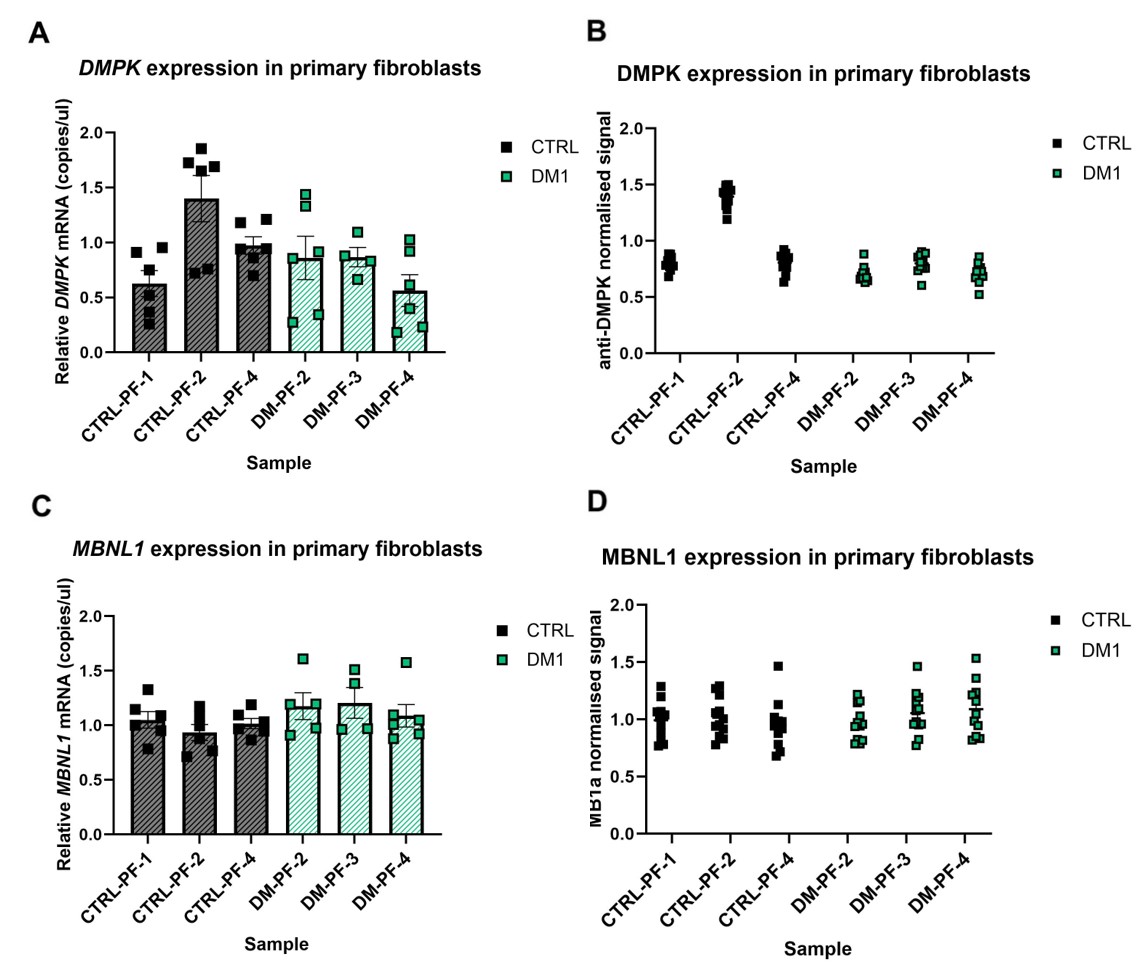
